# Metformin promotes broad neuroprotection and proteostatic resilience via rngo/DDI2 stabilisation

**DOI:** 10.64898/2026.02.11.705245

**Authors:** Dongwei Xu, Youngah Kim, Sharifah Anoar, Xinwa Jiang, Argyro Alatza, Riccardo Zenezini Chiozzi, Konstantinos Thalassinos, Adrian M. Isaacs, Tammaryn Lashley, Selina Wray, Teresa Niccoli

**Affiliations:** Department of Genetics, Evolution and Environment, Institute of Healthy Ageing, UCL, Gower Street, London WC1E 6BT, UK; Department of Neurodegenerative Disease, UCL Queen Square Institute of Neurology, UK; Department of Structural and Molecular Biology, UCL, Gower Street, London WC1E 6BT, UK; UK Dementia Research Institute at UCL, Cruciform Building, London WC1E 6BT, UK

## Abstract

Alzheimer’s disease (AD) is one of the most common age-related causes of death, with limited effective disease-modifying treatments. Although metformin shows promise as a disease-modifying agent for AD, its molecular mechanism—specifically how it confers neuroprotection despite potentially increasing amyloid-beta (Aβ) load—remains obscure.

Here, we demonstrate that metformin functions as a pharmacological activator of the ubiquitin-binding protease rngo/DDI2. Through a genetic screen in *Drosophila*, we identified rngo/DDI2 as a potent suppressor of Aβ toxicity. We provide in silico and genetic evidence that metformin interacts with the conserved D257 residue of the rngo/DDI2 RVP domain, inducing homodimerisation and subsequent protein stabilisation. This activation boosts proteasome activity in the presence of Aβ preferentially clearing highly abundant proteins to preserve proteostasis. Crucially, this intervention is broadly effective; rngo/DDI2 upregulation robustly suppresses toxicity in models of TDP-43 and C9orf72-repeat expansion pathology, indicating a generalised mechanism of neuroprotection. Supported by human iPSC data and evidence of DDI2 depletion in AD patient brains, our results identify rngo/DDI2 as a conserved regulator of neuronal resilience. We propose that directly targeting DDI2 stabilisation represents a novel, broadly applicable therapeutic strategy to counteract proteotoxic stress across a broad spectrum of neurodegenerative diseases.

## Introduction

Dementia is the UK’s leading cause of mortality, affecting nearly 1 million people (Office for National Statistics). It remains the only major cause of death without treatment to significantly slow disease progression. The main risk factor is advanced age, and with life expectancy increasing dementia prevalence is expected to grow significantly and pose a formidable burden on healthcare and economy ^1^.

Alzheimer’s disease (AD), the most common type, is characterised by neurodegeneration, intracellular tau tangles, and extracellular amyloid-β (Aβ) plaques derived from APP processing ^2^. While extensive genetic evidence suggests that increasing production of Aβ 1-42 is sufficient to cause aggressive early onset familial forms of the disease ^3–5^, Aβ pathology is also found in 40% of cognitively normal 90-year-olds ^6^. This suggests that while Aβ accumulation increases risk, other factors trigger disease onset.

As the primary risk factor, age likely increases susceptibility to Aβ toxicity. Aged mice ^7^, flies ^8^, and primates ^9^ were found to be more susceptible to Aβ toxicity, and mutant animals with delayed ageing, also showed delayed onset to Aβ toxicity ^10,11^. Treatment with drugs known to slow ageing such as rapamycin ^12,13^ or lithium ^14–16^, can also delay the onset or slow disease in mouse models.

However, benefits do not always translate to humans; for example, rapamycin trials showed increased inflammatory biomarkers ^17^. Because these drugs have broad, pleiotropic effects, pinpointing their specific beneficial mechanisms is crucial for developing targeted therapies.

Metformin, a commonly prescribed Type 2 diabetes (T2D) medication has shown promise as an anti-ageing drug in a number of model organisms ^18–20^, and it improves healthspan over a number of parameters ^21^.

In AD research the results have however been mixed: while some animal studies find metformin worsens Aβ biogenesis and cognitive function ^22–24^, others have shown a protective role ^25–27^, Similarly, epidemiological studies have found both detrimental ^28,29^, and beneficial effects on the risk of dementia ^30^ and small-scale clinical trials indicate metformin can improve cognitive function in early AD patients, without altering Aβ levels ^31,32^, suggesting it modulates cellular resistance to toxicity.

Metformin has broad effects on cellular molecular dynamics: in yeast it modulates the homodimerization of over 700 proteins ^33^. Some of these effects are likely relevant (both positively and negatively) to the development of AD.

In a previous study we established that metformin reduced Aβ toxicity in our fly model without affecting Aβ levels ^34^. To identify the precise targets responsible for this beneficial effect, we screened 128 fly homologues, identifying *rings lost (rngo)*, the homologue of human DDI1/2 (DNA Damage-Inducible 1 Homolog 1/2) as a strong suppressor. We found that metformin binds to a residue in rngo’s conserved protease domain, increasing homodimerisation and protein levels. This boosts proteasomal activity and resistance to stress—a mechanism we confirmed is conserved in human iPSC-derived neurons.

We propose that increased rngo/DDI2 homodimerisation is a promising new therapeutic avenue. Developing drugs that enhance this dimerisation could increase proteasomal activity in dementias, potentially making neurons more resilient to the toxic protein accumulation associated with these diseases.

## Methods

### Bioinformatics identification of homologues

From the list of metformin modulated targets identified in yeast ^33^, the 342 with increased and the 403 with decreased homodimerisation were taken and converted using the DIOPT (DRSC integrative ortholog prediction tool) into *Drosophila* gene homologues. For those with multiple homologues, only the fly genes with the Best Score (Table S1) were taken forward.

### Drosophila stocks

The elavGS stock was derived from the original elavGS-301.2 line ^35^. The w^1118^ (BDSC: 3605) stock was from Bloomington Drosophila Stock Centre (BDSC). da-GS (daughterless-geneswitch) was derived from the original line developed by Monnier’s group ^36^. The UAS-Aβ_42_*2 stock was from Pedro Fernandez Funez (University of Minnesota) ^37^. UASp-rngo^FL^-GFP, UASp-rngo^ΔRVP^-GFP, UASp-rngo^D257A^-GFP lines were kindly provided by Andreas Wodarz (Universität Göttingen). Stocks ordered for the genetic screen were shown in Table S2.

### Lifespan assay

Parental flies were crossed and allowed to lay eggs for 24 hours on agar grape plates with yeast paste. Eggs were collected in PBS and seeded at equal density into bottles with fresh SYA food. 2 days post-eclosion, the female offsprings with the desired genotype were selected into plastic vials containing SYA food (15 flies per vial, 7-10 vials per genotype/condition). Flies were tipped onto fresh food three times per week, and deaths were scored every tipping. Data are presented as cumulative survival curves, and survival rates were compared using log-rank tests. All lifespans were performed at 25°C.

### Climbing assay

Parental flies were allowed to lay eggs in a bottle containing SYA food for 24 hr and then discarded. 2 days post-eclosion, the female offspring with the desired genotype were selected into plastic vials containing SYA food (15 flies per vial, 5 vials per genotype for the genetic screen). The vials were clicked to a Drosoflipper^TM^. Flies were tipped onto fresh food three times per week. Climbing assay was conducted every 2-5 days. Flies were tapped down in front of a webcam, and the climbing process was recorded until the flies reached the top. Videos were analysed in FreeClimber V0.4.0 (github.com/adamspierer/FreeClimber) ^38^ and climbing velocities per vial were calculated, plotted over time and differences were analysed by 2-way ANOVA.

### Bortezomib sensitivity assay

Bortezomib (BTZ, APExBIO A2614) 500 μM stock solution was prepared in EtOH. One day before use, 100 μL of 175 μM BTZ (stock solution diluted in H_2_O) was added on top of SYA food in clean plastic vials free of splash. *da-GS* and *da-GS>rngo-GFP* flies (20 flies each vial, 3 vials each genotype) were kept on RU+ food for 7 days and then placed on RU+ food with BTZ. The flies were tipped three times per week, and deaths were scored every tipping. Data are presented as cumulative survival curves, and survival rates were compared using log-rank tests. All lifespans were performed at 25°C.

### qPCR

Total RNA was extracted from 10-15 fly heads per sample using Trizol (Invitrogen) and subsequently treated with DNAse I (Ambion) for DNA digestion. The RNA was then reverse transcribed using Superscript II (Invitrogen) with oligo(dT) primers. Quantitative gene expression analysis was performed on a 7900HT real-time PCR system (Applied Biosystems) using SYBR-green technology (ABI). Relative quantities of transcripts were determined using the relative standard curve method normalized to eIF1A.

**Table 1.**
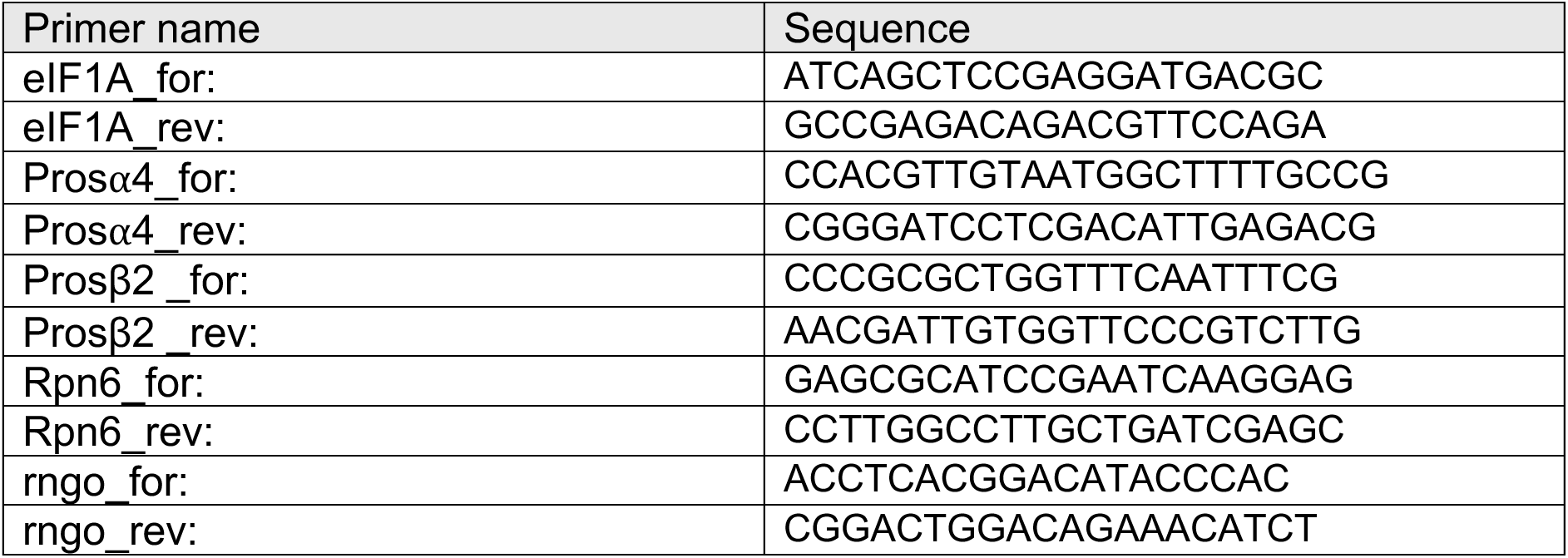
Primers used:

### Protein extraction (*Drosophila*)

Flies were instantly frozen in liquid nitrogen. 10 frozen heads were separated from the bodies and homogenised in 80 µL of 2×LDS sample buffer (Thermo Fisher Scientific), with 50 mM DTT (dithiothreitol) and protease inhibitor cocktail (Roche 11836170001), heated up to 75°C for 5 mins, and spun down at 13000 rpm for 15 mins before loading.

### Protein extraction (human brain)

Brains were donated to the Queen Square Brain Bank (QSBB; UCL Queen Square Institute of Neurology) with full informed consent. Ethical approval for the study was obtained from the NHS research ethics committee and in accordance with the human tissue authority’s code of practice and standards under license no. 12198. Cases underwent standard neuropathological diagnosis according to current consensus criteria. Frozen brain tissue weighing 100 mg was homogenised in RIPA (Sigma R0278) lysis buffer using a Precellys Evolution homogeniser. Subsequently, the homogenates were centrifuged at 4000 rpm for 10 minutes at 4 °C. The resulting homogenised samples, including grey matter and white matter, were subjected to BCA measurement to determine protein concentration. Based on the measured concentrations, diluted samples were prepared to achieve a working concentration of 1 μg/μl. The dilution calculations involved the addition of 25 μl of sample buffer and 10 μl of reducing agent to each 100 μl sample, with the remaining volume supplemented with water.

### Western blot

The samples were loaded onto NuPAGE™ Bis-Tris 4-12% gel (Thermo Fisher Scientific) for electrophoresis. The protein was transferred to Amersham Protran Premium 0.45µm Nitrocellulose or PVDF membrane using Bio-Rad Trans-Blot Turbo Transfer System. After being washed with Milli-Q water, the membrane was stained with Ponceau (Thermo Fisher Scientific) and imaged. Then the membrane was washed in PBST to remove the signal. For detection of Aβ or ubiquitinated proteins, the membrane was heated up in PBS in a microwave until boiling. The membrane was blocked in skimmed milk (5% in PBST) for 1 hour at room temperature. After washing with PBST, the membrane was incubated with primary antibody solution at 4°C with shaking overnight. The next day, the membrane was washed with PBST 3 times (10 mins each), and incubated with secondary antibody solution for 1 hour at room temperature. After being washed 3 times with PBST (10 mins each), the membrane was incubated with ECL (enhanced chemiluminescence) substrate (Thermo Fisher Scientific) for 1 min, imaged and the band intensities were quantified with Cytiva ImageQuant 800.

The antibodies used for western blots: anti-GFP (Merck 11814460001, 1:3000), anti-rngo (a gift from Andreas Wodarz’s group, 1:3000), anti-ubiquitin (Enzo Life Sciences PW8810, 1:4000), anti-actin (Abcam Ab8224, 1:10000), anti-Aβ 6E10 (BioLegend 803001, 1:2000), anti-DDI2 (Santa Cruz sc-514004, 1:1000), anti-mouse (Abcam Ab6789, 1:10000), anti-rabbit (Abcam Ab6721, 1:10000), Lamin A/C (Abcam ab1085951, :1000). Secondaries for the human samples were: goat anti-mouse (Li-Cor;1:10000) Goat anti-rabbit (Li-Cor, 1:10000).

### Chymotrypsin-like proteasome activity assay

The method was adapted from literature ^39^. Flies were anaesthetised with CO_2_ and collected for proteasome activity assay. Either one whole body or seven heads were homogenised in ice-cold 80 µL reaction buffer (Tris-HCl pH 7.5 with 1 mM MgCl_2_, 2 mM ATP, 1 mM DTT). Samples were spun at 13000 rpm, 4°C for 5 mins, and supernatant was added to ice-cold N-Suc-LLVY-AMC (N-Succinyl-Leu-Leu-Val-Tyr-7-Amido-4-Methylcoumarin, Sigma Aldrich S6510) in a black 96-well plate (Corning). The plate was read at excitation 355 nm, emission 460 nm at 25°C every 1 min for 60-75 mins by Tecan Infinite® 200 PRO. Readings for the first 10-15 mins were trimmed. The increase in readings (in AU) per min (the substrate hydrolysis rate) was calculated by linear regression. The substrate hydrolysis rate was normalised to total protein determined by BCA (Thermo Fisher Scientific).

### iPSC neuron differentiation and maintenance

KOLF2.1J iPSCs were cultured in StemFlex (Gibco) in Geltrex matrix (Gibco) coated 6-well plates. For coating, 1 mL of Geltrex (1:1000 in DMEM, Gibco) was added to each well and incubated at 37°C for at least 1 hr. iPSCs were regularly passaged 1:3 ∼ 1:6 with 0.5 mM EDTA (0.5 M EDTA Thermo Fisher, diluted 1:1000 in PBS) when the density reached ∼80%. Cortical neurons were generated based on the protocol by Shi et al. (2012). Briefly, iPSCs were pooled into Geltrex-coated plates and grown to 100% density, and the medium was replaced with induction medium (N2B27 + 1 μM dorsomorphin (Tocris) + 10 μM SB431542 (Tocris)) (day 1). N2B27 formulation: 50% DMEM-F12 GlutaMAX (Thermo Fisher 10565018) + 50% Neurobasal Medium (Thermo Fisher 12348-017) + 0.5 × B-27 supplement (Thermo Fisher 17504-044) + 0.5 × N2 supplement (Thermo Fisher 17502-048) + 1 mM L-glutamine (Thermo Fisher 25030-024) + 0.5 × non-essential amino acids (Thermo Fisher 11140-50) + 50 μM 2-mercaptoethanol (Thermo Fisher 31350-10) + 25U Pen-Strep (Thermo Fisher 15070063) + 25U insulin (Sigma i9278). The induction medium was changed daily. On day 10 post-induction when a neuroepithelial layer had formed, the cells were passaged using dispase, and re-plated as clumps in a 1:50 laminin (Sigma) coated 12-well plate. Afterwards, the cells were cultured in N2B27 medium with half medium change. On day 18 post-induction, the cells were passaged again using dispase, and re-plated in a 1:50 laminin coated 12-well plate. On day 25, the cells were detached with accutase, and replated in a poly-ornithine/poly-D-lysine + 1:100 laminin coated 12-well plate. On day 50, the medium was switched to low-glucose, low-insulin N2B27. The formulation of low-glucose, low-insulin N2B27: 50% DMEM-F12 GlutaMAX (Thermo Fisher 10565018) + 50% Neurobasal-A Medium, no D-glucose, no sodium pyruvate (Thermo Fisher A2477501) + 0.5 × B-27 supplement, minus insulin (Thermo Fisher A1895601) + 0.5 × N2 supplement (Thermo Fisher 17502-048) + 1 mM L-glutamine (Thermo Fisher 25030-024) + 0.5 × non-essential amino acids (Thermo Fisher 11140-50) + 50 μM 2-mercaptoethanol (Thermo Fisher 31350-10) + 25U Pen-Strep (Thermo Fisher 15070063). The cells were maintained until day 72 for drug treatment experiments.

### iPSC neuron drug treatment

On day 72 post-induction, a full medium change was performed. The medium was replaced by fresh low-glucose, low-insulin N2B27 medium with or without 10 mM metformin (in PBS) or 5 µM BTZ (in EtOH). Equal amount of PBS or EtOH was added to the medium as a solvent control. 18 hours later, the cells were harvested for protein extraction and western blots.

### Proteomics

7 day old fly heads were frozen and boiled in lysis buffer (5% sodium dodecyl sulphate (SDS), 5 mm tris(2-carboxyethyl)phosphine (TCEP), 10 mm chloroacetamide (CAA), 100 mm Tris, pH 8.5) for 10 minutes and then sonicated (Q705 Sonicator from Fisherbrand) for 2 min with pulses of 1 s on and 1 s off at 50% amplitude. Protein concentration was estimated by BCA assay (Thermo Fisher Scientific). Protein digestion was automated on a KingFisher APEX robot (Thermo Fisher Scientific) in 96-well format using a protocol from Koenig at al (https://doi.org/10.1016/j.xpro.2023.102536) with minor modifications. The 96-well comb is stored in plate #1, the sample in plate #2 in a final concentration of 70% acetonitrile with magnetic MagReSyn Hydroxyl beads (ReSyn Biosciences) in a protein/bead ratio of 1:2. Washing solutions are in plates #3–5 (95% Acetonitrile (ACN)) and plates #6–7 (70% Ethanol). Plate #8 contained 300 μL digestion solution of 100 mm Tris pH 8.5 and trypsin (Promega) in an enzyme:protein ratio of 1:100. The protein aggregation was carried out in two steps of 1 min mixing at medium mixing speed, followed by a 10 min pause each. The sequential washes were performed in 2.5 min and slow speed, without releasing the beads from the magnet. The digestion was set to 16 h at 37 degrees with slow speed. Protease activity was quenched by acidification with trifluoroacetic acid (TFA) to a final pH of 2 and the resulting peptide mixture was purified on OASIS HLB 96 well plate (Waters). Purified peptides were dried in a Savant DNA120 (Thermo Fisher Scientific)

Dried peptides and phosphopeptides were then dissolved in 0.5% TFA analysed using a Vanquish NEO high-performance liquid chromatography system coupled online to an Orbitrap Eclipse mass spectrometer (Thermo Fisher Scientific). Buffer A consisted of water acidified with 0.1% formic acid, while buffer B was 80% acetonitrile and 20% water with 0.1% formic acid. The peptides were separated by a 1 mm BEH HPLC Column (Waters) and the gradient was 3 to 35% B in 53 min at 50 ul/min. Buffer B was then raised to 55% in 1.5 min and increased to 99% for the cleaning step. Peptides were ionized using the settings from Tsiklauri et al (https://pubs.acs.org/doi/full/10.1021/acs.jproteome.5c00327): spray voltage of 4 kV and a capillary heated at 320°C and vaporizer at 200°C with gas set at 32/5/0. The mass spectrometer was set to acquire full-scan MS spectra (350 to 1400 mass/charge ratio) at a mass resolution of 120K and an automated gain control (AGC) target value of 250% (RF lens of 40% and max injection time of 45ms). For MSMS fragmentation, we chose the DIA approach, with an m/z range 361-1033 m/z, divided into 56 windows (12 m/z each with 1 Da overlap), with an AGC of 1000% and resolution of 15K. All raw files were transformed into htrms format and analysed by Spectronaut v19 with directDIA analysis. We used the library generated automatically using Drosophila proteome (from Uniprot) reference together with MaxQuant contaminants list and standard settings: for normal peptides we used the “BGS factory settings”.

Differential expression analysis was performed in Perseus (v2.1.5.0) using default parameters. Only the proteins with valid values in at least 3 samples were kept. For hierarchical clustering for the heatmap plot, missing values were imputed by normal distribution in Perseus, and the dataset was plotted using ComplexHeatmap package in R.

### Statistical analysis

Statistical analysis was carried out in prism or Excel, unless otherwise stated. Lifespans were compared either by log-rank or cox proportional hazards, as stated. Westerns, qPCRs and chymotrypsin assays are plotted as means with standard deviations, unless otherwise stated and compared by one or two way ANOVAs, followed by a post-hoc comparison as stated.

## Results

### Rngo is a strong suppressor of Aβ toxicity

To identify suppressors of Aβ toxicity acting downstream of metformin, we used a *Drosophila* model of Aβ toxicity where 2 tandem copies of wt Aβ1-42 ^37^ are expressed under the control of the inducible pan-neuronal elavGS driver ^35^. This model is induced post-eclosion by feeding the drug RU486 to limit the expression of Aβ to post-mitotic adult neurons, thus avoiding confounding developmental effects. This model displays Aβ accumulation, which leads to a shortened lifespan, and defects in locomotion, assessed by climbing ability, the innate, reflexive behaviour of flies to ascend when startled ^40^. Similarly to our previous studies ^34^, this model is also rescued by feeding 25 mM of metformin (Fig S1A).

We then used DIOPT (DRSC integrative ortholog prediction tool) to identify *Drosophila* homologues (Table S1) of proteins found to be modulated by metformin in yeast ^33^. From these we selected over-expression lines for the top 41 activated genes and RNAi lines for the top 88 down-regulated genes where lines were available from public repositories (Table S2), and screened for their ability to modulate the climbing defect induced by expression of Aβ in our model (Fig 1).

**Fig 1.**
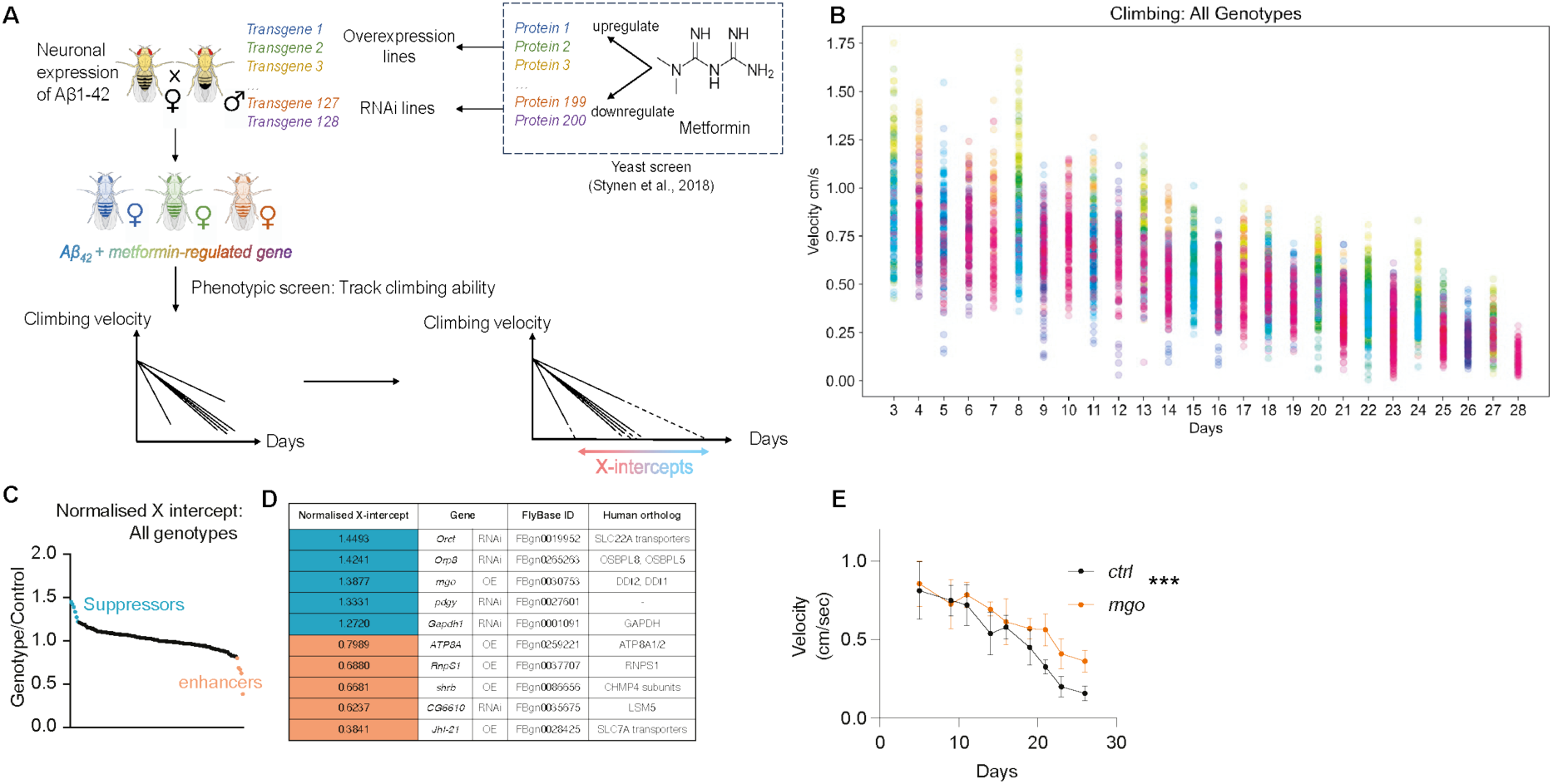
Genetic screen. **A.** Workflow of the genetic screen. The list of metformin-regulated proteins from the published yeast screen were converted into *Drosophila* orthologs by DIOPT. The overexpression and RNAi fly lines were obtained and crossed with the Aβ42 line, resulting in a library of fly lines expressing Aβ42, together with the RNAi or overexpression of a gene. These flies were repeatedly screened for their climbing ability over 25 days, a linear regression for each genotype was calculated and the X-intercepts were calculated for each screened line. **B.** Scatter plot of the climbing velocities of all genotypes in the screen, plotted over time. Each dot stands for a replicate (vial) of 15 flies. Each colour represents one genotype. **C.** Plot of normalised X-intercept values. Candidate suppressors (light blue) and enhancers (light red) of toxicity are highlighted. **D.** A summary of the candidate modifiers of toxicity, the same genotypes are highlighted as in C **E.** Climbing velocity of Aβ-expressing flies with *rngo* overexpression and the control line from the same batch, plotted as means with standard deviations (SD). Significance levels were based on two-way ANOVA ***: *P*<0.001. Genotype: *w*1118; *UAS-Aβ*2/+; elavGS/+, w*1118/*EP-rngo*; *UAS-Aβ*2/*+; *elavGS/*+.

A general workflow is shown in Fig.1A. Each fly line in the screen was crossed to the Aβ model, and 75 female offspring were screened for their climbing performance. We set up the screen in batches of 6-9 lines, always with a control line, which for over-expression stocks was a w^1118^ line and for TRiP (Transgenic RNAi Project) RNAi lines, a v^-^w^+^ control, to match the background of the screened lines. We tracked their climbing velocities three times per week until ∼10% of the flies died and plotted them as a function of age (Fig 1B, Table S3). We applied linear regression to the velocity-age curve, and calculated the X-intercept parameter, which predicts the time at which they stop climbing. We then normalised the X intercepts for each batch against their respective control, to account for potential batch effect (Table S2, Fig 1C). The genotypes with the five highest and five lowest normalised X-intercepts were identified as candidate suppressors and enhancers of Aβ toxicity, respectively (Fig.1C, Table S2). The top candidate suppressors were *Orct*^RNAi^ (*Organic cation transporter*)*, Orp8*^RNAi^ (*Oxysterol-binding protein-related protein 8*)*, rngo* (*rings lost*) over-expression (Fig 1E), *pdgy*^RNAi^ (*pudgy*) and *Gapdh1*^RNAi^ (*glyceraldehyde-3-phosphate dehydrogenase 1*) (Fig S1). The top enhancers were over-expression of *JhI-21* (*Juvenile hormone Inducible-21*), *RnpS1* (*RNA-binding protein S1*) and ATP8 (*ATPase 8A*), together with RNAi of *CG6610* and *shrb* (*shrub*).

Since we were interested in identifying genes responsible for the improvement associated with metformin, we took forward the suppressors. Pdgy, not having a human homologue was dropped. The rest were back-crossed into a w^1118^ (for rngo) or v^-^w^+^ background (for RNAi lines) to remove any potential confounding genetic background effects. To confirm their robustness and identify the most promising suppressors, we used an orthogonal measure of Aβ-toxicity and screened for their ability to rescue the lifespan phenotype of Aβ expressing flies (Fig 2, Fig S1F-H). Only over-expression of *rngo* gave a very slight rescue, which did not reach significance when using a high concentration of the transgene inducer RU486 (200 µM) (Fig 2A). However, when we reduced the RU486 to 50 µM, the Aβ expressing flies still displayed a strong reduction in lifespan (Fig 2B), and neuronal *rngo* over-expression led to a substantial rescue (Fig 2B). Ubiquitous over-expression of *rngo* with 50 µM of RU486 in control flies also led to a lifespan increase (Fig 2C), however over-expression of *rngo* just in neurons during ageing in control flies led to a slight lifespan shortening (Fig S2A, B), suggesting both that the beneficial effects of *rngo* during ageing are mediated by a tissue other than neurons and that the rescue of the Aβ phenotype is specific and not due to a general ameliorating of neuronal ageing. Interestingly, the over-expression of *rngo* had no effect to Aβ levels (Fig 2D), similar to the effect of feeding metformin ^34^. We confirmed by qPCR that our *rngo* over-expression line did indeed lead to over-expression of *rngo* (Fig S2C). Rngo is therefore a robust suppressor of Aβ toxicity and was taken forward for further analysis.

**Fig 2.**
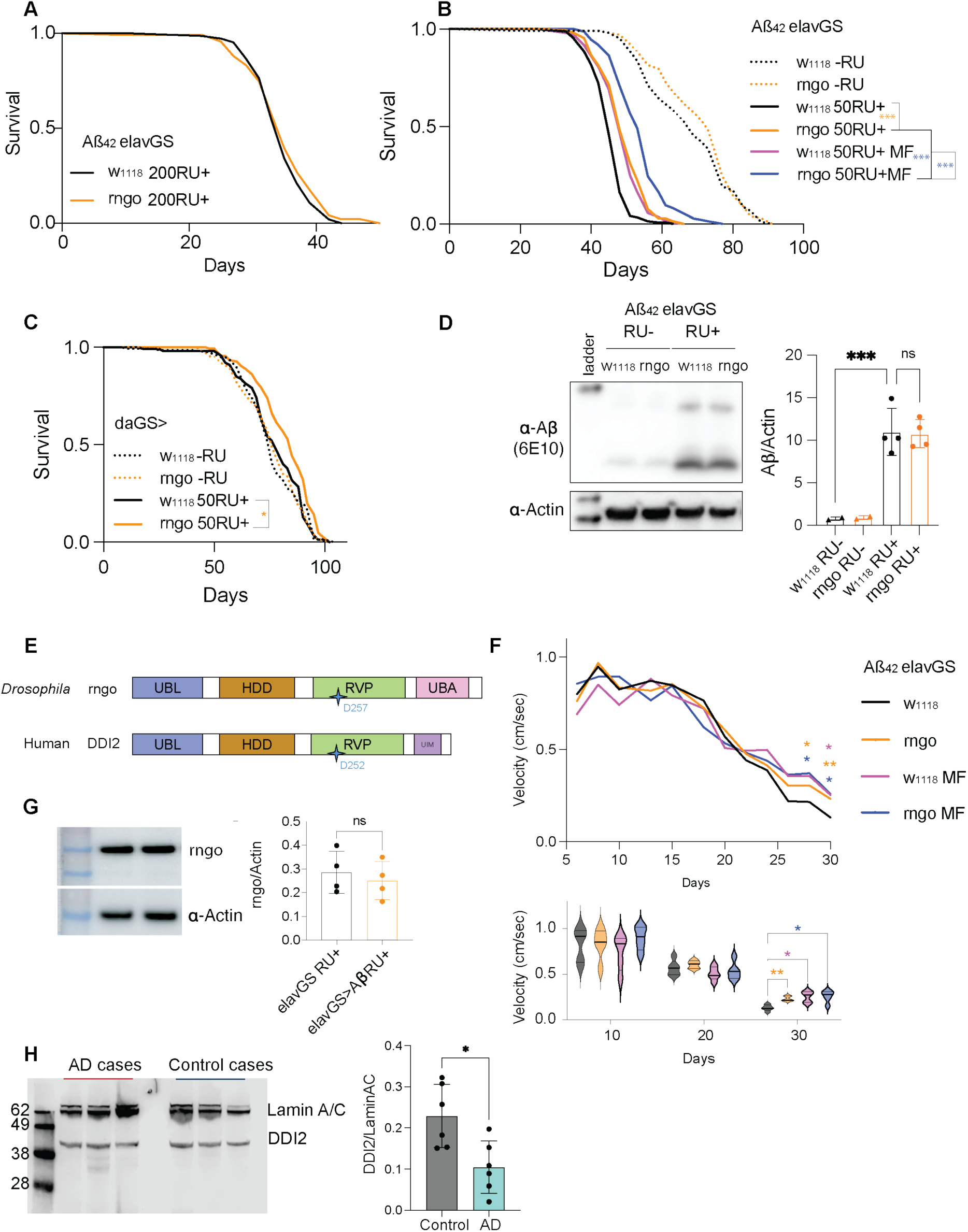
rngo is a robust suppressor of Aβ toxicity. **A.** Lifespan curve of Aβ flies with or without rngo overexpression, using 200 μM RU. Minimum n=137, Comparison by log-rank: ns. Genotype: *w*1118; *UAS-Aβ*2/+; elavGS/+, w*1118/*EP-rngo*; *UAS-Aβ*2/*+; *elavGS/*+. **B.** Lifespan curve of Aβ flies with or without rngo overexpression, with or without metformin (25 mM) feeding. 50 μM RU was added to the food. RU-conditions (dotted lines): minimum n=104, RU+ conditions: Minimum n=146, comparison by log-rank ***p<0.001. Genotype: *w*1118; *UAS-Aβ*2/+; elavGS/+, w*1118/*EP-rngo*; *UAS-Aβ*2/*+; *elavGS/*+. **C.** Lifespan curves of flies expressing rngo in the whole fly, induced with 50 µM RU. Minimum n=145, Comparison by log-rank *p<0.01. Genotype: *w*1118; *da-GS/+, w*1118/*EP-rngo*; *da-GS/*+. **D.** Western blot showing Aβ levels (detected by 6E10 antibody) in fly heads (day 11) with the Aβ transgene uninduced (RU-) or induced (RU+). Actin was used as the loading control for quantification. Quantifications are plotted as means with SD and significance levels were based on two-way ANOVA with Fisher’s LSD test ***p<0.001, RU+ conditions: n=4, RU-conditions: n=2, with 10 heads per replicate. Genotype: *w*1118; *UAS-Aβ*2/+; elavGS/+, w*1118/*EP-rngo*; *UAS-Aβ*2/*+; *elavGS/*+. **E.** Schematic overview of the domains of rngo and DDI2. UBL: Ubiquitin-like. HDD: Helical domain of Ddi1. RVP: Retroviral protease-like. UBA: Ubiquitin associated. UIM: Ubiquitin interacting motif. Star (D257 on rngo, D252 on DDI2) indicates the catalytic centre**. F.** Plot over time of climbing velocities of Aβ-expressing flies with or without rngo co-expression (200 µM RU) and with or without metformin (MF) treatment (above) and violin plot for velocities at 3 timepoints (below). n=5 vials, with 15 flies each vial. Significance levels were based on Dunnett’s multiple comparisons (using w1118 0 mM as the baseline). *p<0.05, **p<0.01. Genotype: *w*1118; *UAS-Aβ*2/+; elavGS/+, w*1118/*EP-rngo*; *UAS-Aβ*2/*+; *elavGS/*+. **G**. Western blot for rngo on 27 day old flies Aß expressing flies and driver alone controls, normalised to actin. p=ns by t-test. **H**. Western blot for DDI2 on of cortical samples from AD cases and controls, relative to LaminAC. The quantification is from 2 gels of different samples. *p= 0.0127 by t-test.

Rngo is a highly conserved ubiquitin shuttling factor belonging to the family of Ddi1-like family proteins, named after the yeast protein Ddi1 (DNA damage inducible 1) ^41^. In humans there are two homologues, DDI1 and 2, where DDI1 is testis-specific, whereas DDI2 is ubiquitously expressed (Fig 2E). Ddi1-like proteins harbour a conserved aspartic protease domain (RVP) that resembles the retroviral aspartyl protease from HIV (human immunodeficiency virus). The UBA (ubiquitin associated) domain, or UIM (ubiquitin-interacting motif) in higher organisms, binds to ubiquitinated proteins, and the UBL (ubiquitin-like) domain binds to proteosomes, thus bringing the ubiquitinated proteins to proteosomes for degradation ^42^. The UBL domain can also bind to ubiquitin with strong affinity ^43^. Ddi1 is the only identified protein that has both ubiquitin-binding ability and protein-cleaving activity other than the 26S proteasomes ^44^.

We asked whether metformin and rngo over-expression act in the same pathway, by treating rngo over-expressing flies with metformin and monitoring their climbing ability and lifespan (Fig 2B, F). We found that rngo over-expression and metformin feeding ameliorated both lifespan and climbing, as expected, and when metformin was fed to rngo over-expressing flies (Fig 2F) there was no further increase in climbing, suggesting the two interventions act in the same pathway. However, the two combined interventions led to an additive extension of lifespan (Fig 2B), suggesting that metformin exerts further beneficial effects on lifespan, maybe acting via other tissues, or via other lifespan-specific pathways.

### Human rngo homologue DDI2 is reduced in Alzheimer’s disease brain

It was recently reported that DDI2 association with the proteasome decreases in AD patient cortical tissue ^45^. To check whether rngo/DDI2 itself was modulated by disease, we checked whether the expression of rngo/DDI2 was changed in our AD fly model and in patient cortical samples. Although rngo was not affected by Aβ expression in flies (Fig 2G), DDI2 levels are decreased in the cortex of AD patients relative to age-matched controls (Fig 2H). Together, this shows that DDI2 is expressed in cortical tissue and its expression and activity are impaired in AD brains, suggesting direct human-relevance and therapeutic potential of our findings in *Drosophila*.

### Metformin binds and stabilises rngo

We next sought to understand how metformin influences rngo protein. We checked whether metformin increased the levels of rngo, but found that metformin did not increase rngo mRNA (Fig 3A) or protein levels (Fig 3B) in fly heads expressing Aβ.

**Fig 3.**
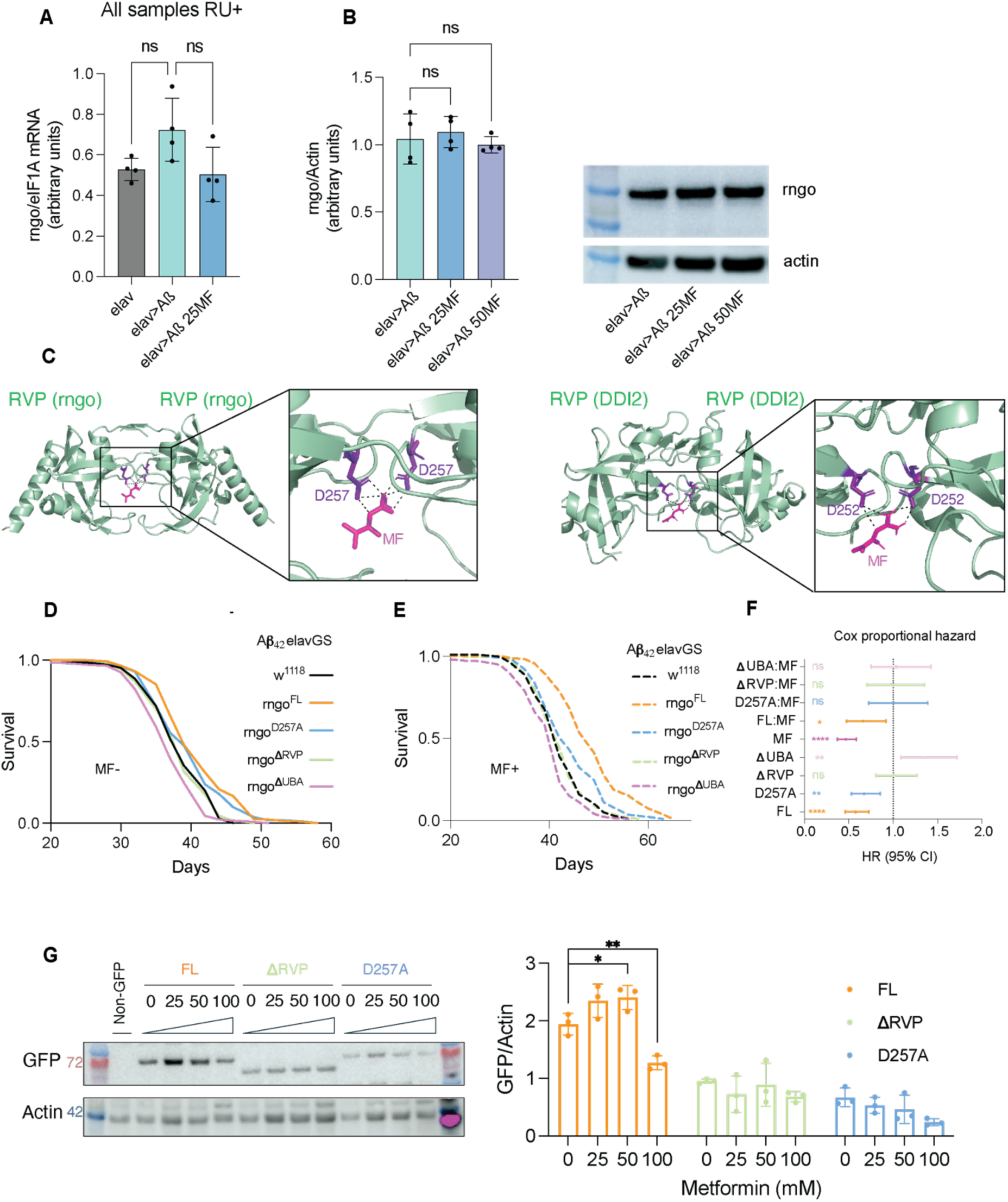
Metformin rescue depends on D257 in the RVP domain. **A**. mRNA levels of rngo in fly heads measured by qPCR and normalised by eIF1A, compared by one way ANOVA. Genotype: *w*1118; *+/+; elavGS/+*, *w*1118; *UAS-Aβ*2/+; elavGS/+.* **B**. Western blot and quantification of rngo protein levels in 7 day old fly heads, normalised by actin, compared by one way ANOVA. Genotype: *w*1118; *UAS-Aβ*2/+; elavGS/+.* **C**. Left: Alphafold3-predicted structure of the rngo RVP dimer, and its putative interaction with metformin. Right: Swissdock prediction of metformin’s interaction with the DDI2 RVP dimer (PDB: 4RGH). Metformin is shown in magenta. **D-E**. Lifespan curves of Aβ-expressing flies with (**E**) or without (**D**) the feeding of 25 mM metformin (shown as 0 mM or 25 mM MF in the legend), with or without overexpression of the full-length (FL), D257A, ΔRVP, or ΔUBA forms of rngo. Minimum n=148. Significance levels were based on log rank t tests. Comparisons were made to Ctrl 0 mM (for all conditions not fed with metformin), or to Ctrl 25 mM MF (for all conditions fed with metformin). Genotype: *UAS-Aβ*2*/+; *elavGS*/+, *UAS-Aβ*2/+*; *elavGS/UASp-rngo-GFP*, *UAS-Aβ*2*/+; *elavGS/UASp-rngo^D257A^-GFP*, *UAS-Aβ*2*/+; *elavGS*/*UASp-rngo^ΔRVP^-GFP*, *UAS-Aβ*2*/+; *elavGS*/*UASp-rngo^ΔUBA^-GFP*. **F**. Cox proportional hazard analysis results showing the hazard ratios (HR) with the 95% confidence interval (CI). Significance levels were based on Wald test. **G**. Western blot results showing the level of wild type and mutant rngo-GFP in the heads of day 9 Aβ flies feeding different metformin doses (in mM). Significance levels were based on Dunnett’s tests following two-way ANOVA. Genotype: *UAS-Aβ*2*/+; *elavGS*/+, *UAS-Aβ*2/+*; *elavGS/UASp-rngo-GFP*, *UAS-Aβ*2*/+; *elavGS/UASp-rngo^D257A^-GFP*, *UAS-Aβ*2*/+; *elavGS*/*UASp-rngo^ΔRVP^-GFP*, *UAS-Aβ*2*/+. ns: P>0.05, *: P<0.05, **: P<0.01, ***: P<0.001.

Given the lack of modulation of rngo levels, we hypothesised that metformin was enhancing rngo function, possibly by direct binding. To understand whether metformin was binding to rngo directly we used *in silico* modelling of their interaction using Alphafold 3, and found that metformin was predicted to bind to D257 on rngo and stabilise a rngo dimer conformation (Fig 3C), in line with the yeast study that reports increased DDI1 homodimerisation levels following metformin treatment ^33^. The structure of RVP domain of human DDI2 has been resolved (PDB: 4RGH), we used Swissdock to dock metformin to the domain and found that metformin was predicted to bind to the equivalent, conserved aspartate D252 residues of DDI2 (Fig 3C).

To confirm the physiological relevance of this interaction, we over-expressed GFP-tagged *rngo* constructs lacking the RVP or the UBA domain and a D257A mutant, in Ab expressing flies ^46^. If the D257 residue was essential for rngo’s interaction with metformin, mutants lacking that residue (D257A and RVPΔ) would abolish the rngo interaction with metformin. First, we established the baseline effects of the proteins alone. Full-length (FL) *rngo* significantly extended lifespan, reducing the hazard ratio (Fig 3 D, F), as expected, whereas *rngo* lacking the RVP domain had no effect, and a UBA deletion proved deleterious. The D257A mutant, which lacks protease function, conferred a minor lifespan extension. This suggests that while the protease function is partially dispensable for rescue, the RVP domain is essential (Fig 3D, F). We then fed metformin to all our lines. While metformin extended the lifespan of all groups (likely via endogenous *rngo*), it showed a significant statistical interaction only with the FL *rngo* over-expression (Fig 3E, F). In this group, the combined lifespan extension was synergistic, greater than the sum of the individual effects of the drug and the FL rngo (Fig 3E, F), likely because metformin increased the levels of full length GFP tagged rngo (Fig 3G), unlike what we found with endogenous rngo (Fig 3B). This discrepancy could be potentially due to a feed-back inhibition mechanism that limits the increase of endogenous rngo mRNA and protein levels. However, when the protein is expressed from an exogenous promoter, as the rngo-GFP, it might escape such strong feedback inhibition.

Conversely, this synergy was lost in the mutant lines, where the interaction term in the hazard analysis was not significant. This implies that both the RVP domain and the D257 residue are mechanistically required for metformin to potentiate rngo function *in vivo*.

Overall, these results suggest that metformin binds to rngo/DDI2 directly, likely via D257/D252, to stabilise it as a homodimer, and increase its activity.

### Metformin, via rngo, enhances resistance against proteasomal stress

Human DDI2 plays a critical role in maintaining cellular proteostasis during proteasomal stresses. Aβ is known to inhibit the proteasome ^47^, leading to increased aggregation of poly-ubiquitinated proteins and proteotoxic stress in neurons ^34^. We therefore investigated whether metformin’s rescue via rngo was mediated by its ability to alleviate proteasomal stress. Both the overexpression of DDI2 and metformin treatment in cancer cell lines have been shown to be sufficient to confer resistance against bortezomib (BTZ), a proteasomal inhibitor ^48,49^. To examine whether metformin and rngo could enhance flies’ resistance against proteasome inhibition *in vivo*, we over-expressed rngo-GFP ubiquitously, to circumvent the low BBB permeability of BTZ ^50^. We induced rngo-GFP for 7 days and then treated the flies with metformin, and BTZ for 24 hours, we found that metformin treatment alone or the overexpression of rngo alone was sufficient to suppress the accumulation of ubiquitinated proteins in BTZ-treated flies (Fig 4 A,B), while rngo overexpression together with metformin treatment did not further reduce ubiquitinated proteins (Fig 4 A,B). Over-expression of *rngo* also increased survival on food with BTZ (Fig 4 C).

**Fig 4.**
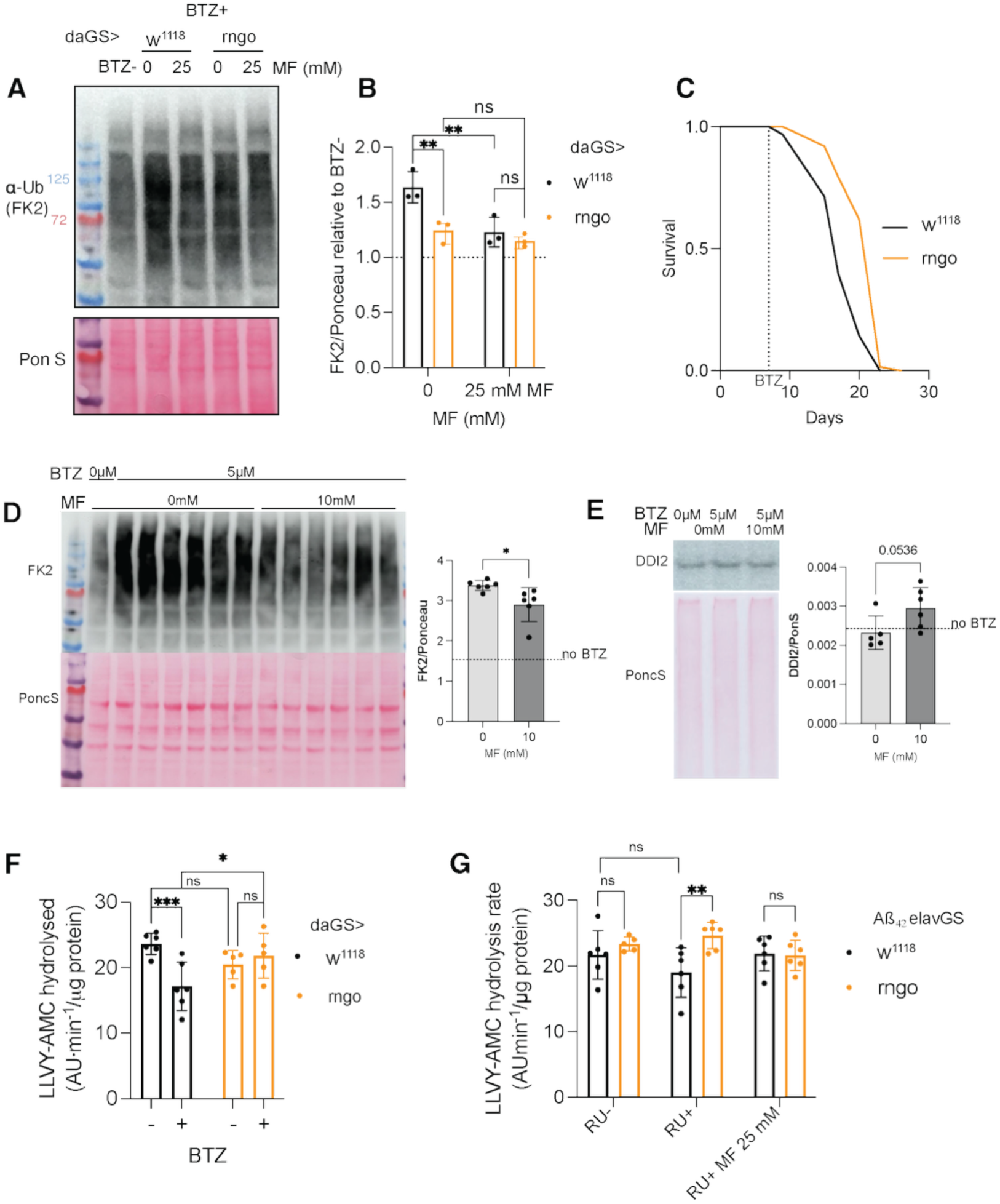
Metformin and rngo enhance resistance against proteasome inhibition. **A-B.** Western blot results showing ubiquitinated proteins (detected by the FK2 clone antibody) in flies with or without rngo over-expression, with or without feeding 25 mM metformin. Ponceau staining was used as the loading control. Significance levels were based on Fisher’s LSD test following two-way ANOVA. Genotype: *da-GS*/+; +/+, *da-GS*/+; *UASp-rngo-GFP*/+. **C.** Survival curve of control flies and rngo-overexpressing flies feeding BTZ-containing food from day 7. Minimum n=63. log rank t test: P<0.001. Genotype: *da-GS*/+; +/+, *da-GS*/+; *UASp-rngo-GFP*/+. **D**. Western blot showing ubiquitinated protein signals (detected by FK2 antibody) in iPSC-derived cortical neurons untreated or treated with BTZ (5 μM) with or without metformin (10 mM) co-treatment for 18 hours. Ponceau staining was used as the loading control. n=6 wells of cells. Significance levels were based on unpaired t tests. **E**. Western blot showing the level of DDI2 in day 72 iPSC-derived neurons treated with BTZ and meformin. n=5 or 6 wells of cells. Significance levels were based on unpaired t tests. **F.** Chymotrypsin-like proteasome activity (measured by the hydrolysis rate of LLVY-AMC substrate and normalised by total protein) in flies with or without rngo-overexpression, with or without Bortezomib treatment. n=6 for control, n=5 for rngo. Significance levels were based on Fisher’s LSD test following two-way ANOVA. Genotype: *da-GS*/+; +/+, *da-GS*/+; *UASp-rngo-GFP*/+. **G.** Chymotrypsin-like proteasome activity (measured by the hydrolysis rate of LLVY-AMC substrate and normalised by total protein) in flies with or without rngo-overexpression, with or without Bortezomib treatment. n=6 for control, n=5 for rngo. Significance levels were based on Fisher’s LSD test following two-way ANOVA. Genotype: *da-GS*/+; +/+, *da-GS*/+; *UASp-rngo-GFP*/+. **G.** Effect of Aβ, rngo co-expression, and metformin on the chymotrypsin-like proteasome activity. n=6, with 7 heads per sample. Significance levels were based on Tukey’s tests following two-way ANOVA. Genotype: *UAS-Aβ*2*; *elavGS*, *UAS-Aβ*2*; *elavGS*/*UASp-rngo-GFP*. ns: p>0.05, *: p<0.05, **: p<0.01, ***: p<0.001.

We then checked whether the beneficial effects of metformin on protein homeostasis are conserved in human neurons. iPSC derived cortical neurons were treated with 5 µM BTZ to induce proteasomal stress, either with or without 10 mM metformin co-treatment. We found that metformin led to a clearance of poly-ubiquitin accumulation induced by BTZ treatment (Fig 4D), suggesting that the protective effect of metformin on proteasomal stress is conserved. We confirmed DDI2 was expressed in cortical neurons, with the metformin treatment of BTZ exposed cells leading to a slight, non significant increase in DDI2 (Fig 4E).

DDI2 is known to act on the proteasome to boost its activities ^51^. To check whether the increased resistance to BTZ was due to a boost in proteasome activity we measured chymotrypsin-like proteasome activity and found that whereas BTZ treatment was able to reduce proteasome activity as expected, rngo over-expression protected the proteasome from BTZ inhibition (Fig 4F), suggesting the increase in clearance of poly-ubiquitinated proteins by rngo over-expression is due to increased proteasomal activity.

We next checked whether this was also the case in Aβ over-expressing brains and found that expression of Aβ showed a trend towards inhibition of the proteasome, consistent with what has been found in AD models and patient tissue ^45^, although this did not reach significance in a post-hoc comparison, likely due to the variability of the assay. Expression of rngo boosted the chymotrypsin-like activity of the proteasome when Aβ was expressed (+RU) (Fig 4G). In the presence of metformin, rngo was not able to boost the proteasome any further, suggesting metformin and rngo act in the same pathway (Fig 4G). In addition, rngo and metformin decreased poly-ubiquitinated protein levels in Aβ over-expressing brains (Fig S3).

Mammalian DDI2 is known to boost the proteasome in multiple ways, it can bind directly to the proteasome, delivering poly-ubiquitinated for degradation ^52^ and it can cleave and activate the nuclear translocation of NFE2L1 (Nuclear factor erythroid 2-like 1), a highly conserved transcription factor regulating proteasome biogenesis ^53–55^. To check whether rngo acted via cnc-C (the fly homologue of NFE2L1), we monitored levels of cnc-C in the nucleus in response to rngo over-expression and found that there was no change in cnc-C nuclear levels (Fig S4). NFE2L1/cnc-C is regulated by a complex feedback mechanism ^56^, to confirm there was no activation of cnc-C we therefore also checked downstream, at the mRNA levels of proteasomal subunits modulated by cnc-C: *Prosβ2*, *Prosα4* and *Rpn6* (Fig S4) and found none were increased in rngo over-expressing or metformin-treated flies (Fig S4), suggesting the rngo rescue is not via cnc-C.

Together these results indicate that the over-expression of rngo, downstream of metformin treatment, is able to protect the proteasome from Aβ toxicity, and boost protein homeostasis, independently of cnc-C. It is therefore possible that rngo is boosting proteasomal activity through its described shuttling activity ^52^.

### Aβ expression decreases proteasomal subunits

To understand how rngo over-expression was modulating the proteome of Aβ expressing flies as a whole, we carried out untargeted proteomic analyses on fly heads expressing Aβ with and without rngo over-expression, and a driver alone control. We identified over 4400 proteins and PCA analysis showed separation of the samples (Fig S5), suggesting good quality data. We found that Aβ expression, relative to the driver only control, was leading to a substantial change in protein levels, with 1450 proteins differentially expressed (DE). GO enrichment analysis showed that upregulated proteins were strongly associated with pathways linked to mitochondrial oxidative phosphorylation (Fig 5A, Table S4), as has been seen in mouse models of disease ^57^. Downregulated proteins, on the other hand, include those associated with cytoplasmic translation (Fig 5B, Table S4), in keeping with what has been found in AD patients and models ^58^, and, interestingly, the proteasome, with 31 subunits showing significant down-regulation (Fig 5B, Table S4). This is consistent with the drop in translation and the proteasomal dysfunction seen in patients and models of disease ^45,59^ and suggests clear impairment in protein homeostasis in response to Aβ expression.

**Fig 5.**
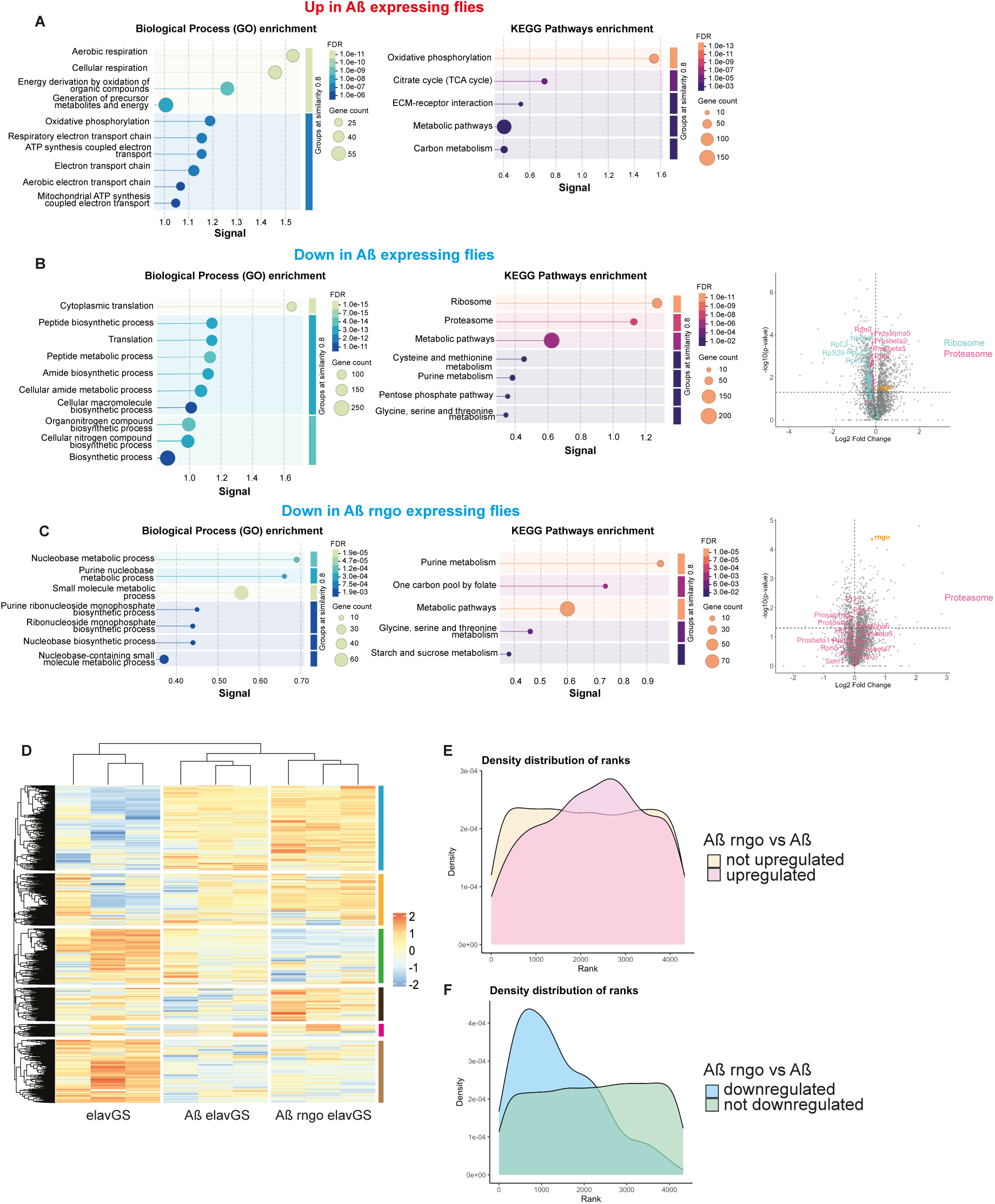
Proteomic analysis of rngo over-expressing heads. **A.** Most enriched GO categories in proteins up-regulated in Aβ expressing fly heads. **B.** Most enriched GO categories in proteins down-regulated in Aβ expressing fly heads, and volcano plot for the comparison of Aβ vs driver alone control showing ribosomal and proteasomal proteins.**C.** Most enriched GO categories in proteins down-regulated when rngo is over-expressed in Aβ expressing fly heads, and volcano plot for the comparison of Aβrngo vs Aβ alone showing proteasomal proteins. **D.** Hierarchical clustering of proteins. The values were standardised across each row independently, and the *z*-scores were plotted. **E.** Density plot showing the distribution of the rank of proteins (by intensity) up-regulated or **F.** down-regulated by rngo overexpression. Genotype: *w*1118; *UAS-Aβ*2; elavGS, EP-rngo*; *UAS-Aβ*2*; *elavGS*.

### Rngo over-expression clears abundant proteins

Over-expression of rngo leads to changes in 700 proteins and interestingly, hierarchical clustering of protein levels across all samples (Fig 5D) indicates that only one cluster (black), associated with mitochondrial translation shows a normalisation back to control level (Fig S6). Overall, however rngo does not normalise protein concentrations back to control levels, indicating the rescue is potentially mediated by the activation of compensatory measures.

Enrichment analysis of differentially expressed proteins shows that rngo did not lead to over-expression of proteins associated with any particular pathways and down-regulated proteins were weakly associated with nucleobase and purine metabolism (Fig 5C), suggesting that the rescue is not due to strong changes in any given molecular pathway. A volcano plot showed that the proteasome subunits were not upregulated in response to rngo expression (Fig 5C), and therefore the increase in proteasome activity we observed can’t be due to an increase in proteasome complexes.

Given the changes in proteasomal activity we looked in more detail as to whether the proteins modulated in Aβ and following rngo over-expression were particularly enriched for proteins targeted to the proteasome. We looked, amongst our DE protein list, for proteins predicted to be targeted to proteasome, autophagy or endosomal microautophagy (containing KFERQ sequences), as described in literature ^60^. We found that proteins reduced in Aβ expressing flies were enriched for substrates of both autophagy and the proteasome (Fig S7), suggesting that the proteasomal dysfunction is accompanied by autophagy dysfunction too, as has been described in AD models and patients ^61^ ^62^.

There was no enrichment for any particular class of substrates upon rngo over-expression suggesting that the rescue is not associated with a particular degradation pathway.

It has been suggested that neurogenerative disease, in particular AD, are associated with aggregation of abundant proteins at the limit of their solubility ^63^, we therefore asked whether rngo, with its ability to boost the proteasome, might be selectively degrading abundant proteins. We ranked proteins from the most to the least abundant protein, and indeed found that abundant proteins were selectively decreased by rngo (Fig 5E,F). This data suggests that over-expression of rngo leads to the preferential reduction of highly abundant proteins, which might in turn allow the cell to deal with the proteotoxic stress caused by Aβ over-expression.

### Rngo rescues a number of neurodegenerative models

Proteasomal impairments^64^ and the increased aggregation of proteins at the edge of solubility ^63^ has been proposed as a common toxicity mechanism across a number of neurodegenerative diseases. We asked therefore whether rngo over-expression could rescue other models of neurodegeneration. We used two models of frontotemporal dementia and amyotrophic lateral sclerosis expressing the most common pathological hallmarks in these diseases: TDP-43 ^65^ and the dipeptide repeat proteins (DPRs) produced by a G4C2 repeat expansion found in the gene C9orf72 (C9) ^66^. Expression of human TDP-43^Q331K^, (G4C2)_36_ pure repeats and GA repeats, which are the most common DPRs found in patient brains ^67^, lead to a shortened lifespan and neuronal toxicity in flies ^68,69^ (Fig 6). Increased rngo expression in all these models extended their shortened lifespan (Fig 6), suggesting that enhanced rngo activity might be a viable therapeutic option across neurogenerative models.

**Fig 6.**
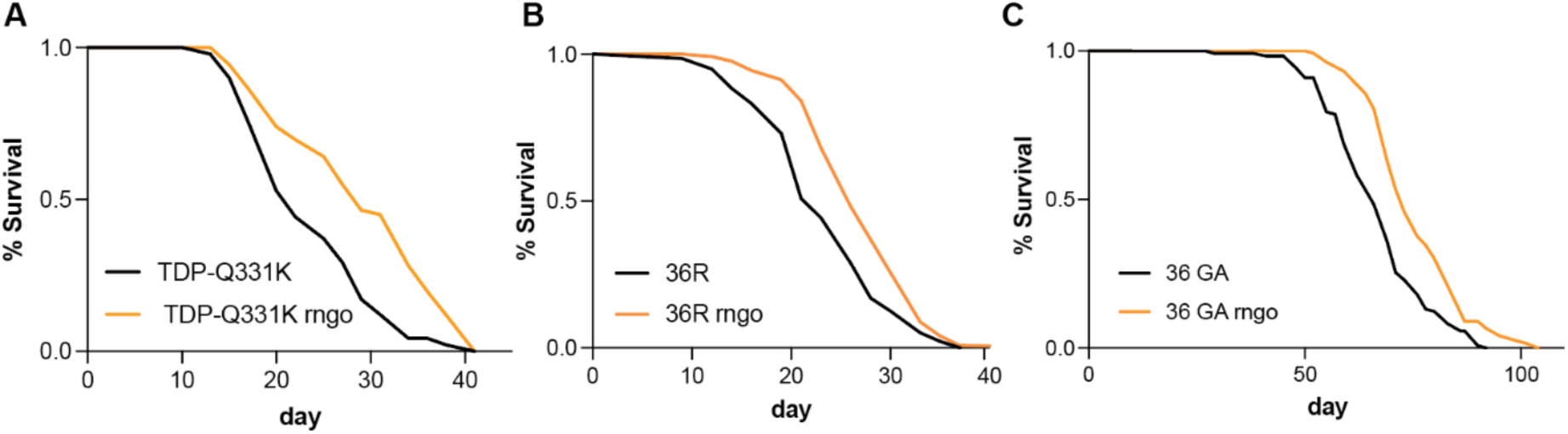
Rngo increases lifespans of ALS/FTD models. Lifespan of **A.**TDP^Q331K^, **B.** 36R and **C.** 36GA expressing flies with (orange) and without rngo over-expression (black). The GA lifespan was run on 50µM RU (n=122-150), conditions compared by log-rank, p= 1.54388E-09 for TDP^Q33K^, p= 3.85722E-05 for 36R and p= 1.75614E-07 for GA). Genotypes: *w1118*; *UAS-TDP^Q331K^; elavGS, w1118/EPrngo; UAS-TDP^Q331K^; elavGS, w1118;UAS-36R; elavGS, w1118/EPrngo; UAS-36R; elavGS, UAS-36GA; elavGS, w1118/EPrngo; UAS-36GA; elavGS*

## Conclusions

Metformin has shown some beneficial effects on Alzheimer’s disease in patient studies ^31,32^, however its effects are not consistent across studies. This variability is likely because, in addition to its benefits, metformin has also been shown to increase APP processing, potentially leading to Aβ accumulation ^22^.

Taking advantage of the strong suppression of Aβ toxicity by metformin in a fly model ^34^, we used this system to pinpoint how the drug’s beneficial effects are mediated. We conducted a large-scale genetic screen on known metformin targets and identified rngo/DDI2, a highly conserved ubiquitin-binding protease, as a mediator of metformin’s protective effect against Aβ toxicity. Importantly, we show that rngo over-expression can rescue a number of different models of dementia, suggesting its beneficial effects might be broadly relevant to a range of neurodegenerative diseases.

Our data suggests that metformin directly binds to rngo/DDI2 to increase its dimerisation and stability. This, in turn, leads to increased proteasomal activity, which is beneficial in the context of toxicity linked to Aβ and other neurodegeneration-associated insults.

While the exact mechanism for the increased proteasomal activity is still unclear, DDI2 is known to cleave and activate NFE2L1 ^70^, a transcription factor regulating proteasomal subunit expression. Furthermore, a recent study suggested that metformin’s beneficial effects on cynomolgus monkey brain aging are at least partially mediated by NFE2L2 activation ^71^. Drosophila has a single homologue of both NFE2L1 and 2: cnc-C, yet we could not see any cnc-C activation following rngo over-expression, and a rngo mutant predicted to have no proteolytic activity still rescued toxicity, suggesting that it is unlikely to be a key mediator of the beneficial effect of metformin on Aβ toxicity.

DDI2 has been shown to directly bind the proteasome and ubiquitinated proteins, acting as a shuttling factor that targets them for degradation ^52^. Rngo, like its yeast homologue Ddi1, has a UBL and a UBA domain. The UBL domain of the yeast and human homologues ^52,72,73^ binds the proteasome, and the fly UBA domain binds ubiquitin ^46^. We have found that the over-expression of the RVP deleted rngo does not have a phenotype, presumably because it can’t dimerise with the endogenous rngo and interfere with its function. However, UBA deleted rngo acted as a dominant negative, likely because it can dimerise with the endogenous rngo and then interfere with its ability to shuttle ubiquitinated proteins to the proteasome.

Other than Ho and multiple K48-ubiquitinated proteins in yeast ^74,75^, specific clients of the Ddi1-like family have not been extensively characterised. While it is possible the rescue is due to the specific degradation of particular clients, our proteomic analysis revealed that highly expressed proteins were reduced in rngo over-expressing flies. This suggests that the degradation of a wide range of highly available proteins, rather than specific clients, might be facilitating the rescue.

Our proteomic analysis revealed that Aβ expression causes a significant reduction in both ribosomal and proteasomal proteins, strongly suggesting a major impairment of protein homeostasis. This pattern is similar to what has been observed in AD patient brains ^76^. Concurrently, we observed an increase in mitochondrial proteins. This specific pattern is highly relevant because it has also been observed in the aging brain ^77^, possibly indicating that aging-associated impairments might predispose to AD. The decreased availability of ribosomal proteins, and the ensuing reduction of translating ribosomes, has been suggested to favour the translation of mRNAs with a high translation initiation affinity, such as a subset of respiratory chain components^77^. Under conditions of low ribosome availability, these high-affinity transcripts are thought to benefit through two mechanisms: first, they out-compete weakly translated mRNAs for the limited ribosomal pool ^78^ and secondly, the lower total number of ribosomes on them relieves ribosomal trafficking and pausing events, thereby increasing the protein synthesis rate of the corresponding proteins ^77^, which in turn leads to the observed increase in mitochondrial proteins.

Overall, we propose that Aβ expression in neurons leads to severe proteomic homeostasis impairments, characterized by a significant drop in both translation (ribosomes) and protein turnover (proteasomes). This dual impairment affects both the production and clearance of neuronal proteins, aligning with observations in post-mortem patient brains ^45^. In this compromised context, the clearance of highly abundant proteins observed downstream of rngo over-expression may act to relieve cellular stress and increase neuronal resilience to the toxic effects of Aβ.

Highlighting its clinical relevance, we show that DDI2 is reduced in patient brain samples, and a recent analysis has shown that in AD patient brains there is reduced binding of DDI2 to the proteasome ^45^.

Our study therefore not only helps explain the mechanism by which metformin achieves a rescue of Aβ toxicity, but it also identifies a therapeutically relevant target in vivo. Specifically, we pinpoint a clear therapeutic avenue: enhancing rngo/DDI2 function by targeting its RVP domain to increase its dimerisation provides a viable strategy for therapeutic intervention in neurodegeneration.

## Acknowledgements

Post-mortem human brain tissue was received from The Queen Square Brain Bank is supported by the Reta Lila Weston Institute of Neurological Studies, UCL Queen Square Institute of Neurology. We thank Nazif Alic for helpful discussions and comments on the manuscript.

